# Neural Modulation of Social Distance on Third-Party Punishment

**DOI:** 10.1101/274720

**Authors:** Chen Qu, Zixuan Tang, Huijun Zhang, Yang Hu, Jean Claude Dreher

## Abstract

As a crucial mechanism to enforce social norms, people as third parties tend to punish the norm violators even it costs their own pay-off. However, people do not usually treat everyone equally, e.g., it is shown that people are nice to close others. Here, we investigated how third party punishment (TPP) and its neural correlates is modulated by social distance (SD) by using fMRI. Behaviorally, participants punished more when the unfair perpetrator was more distant to them. Such SD-modulation effect was stronger when the punishment was free. Model-based results showed that SD-dependent computational signals were encoded in right dlPFC. More interestingly, SD modulated the relationship between punishment levels and neural activities in default network including vmPFC and bilateral hippocampus. The explorative functional connectivity analysis further showed that the vmPFC increased the association with left dlPFC when participants punished close others. Finally, punishment type (costly vs. free) also modulated the relationship between punishment levels and neural correlates in dACC and the ventral striatum. Taken together, our results revealed the neurocomputational underpinnings of how SD plays an important role in affecting TPP.

## 1. Introduction

Social norm is of great importantce in human society as it maintained the social coherence and social cooperation. The ability to develop norms and enforce them through the use of sanctions is thought to be one of the distinguishing characteristics of the human species (Boyd & Richerson, 1988; Ernst Fehr & Fischbacher, 2003). The sanction may be performed either by opponent or by impartial bystanders, the former means the sanction is taken by individuals whose economic payoff is directly harmed by the norm violation, while the latter means the sanction is taken by “third parties”, who are unaffected by the deviation but in a position to punish the violator (Bendor & Swistak, 2001; Ernst Fehr & Fischbacher, 2004; Ostrom, 2000). The third party punishment task including a proposer, a receiver, and an observer. The proposer and the receiver had an initial endowment, the proposer can decide the allocation while the receiver could only accept the proposal. Participants as an observer could punish the proposer by their own payoff. As the sanction by opponent is observed in multiple social species, third party punishment has only be seen in human species (Ernst Fehr & Fischbacher, 2004; Riedl, Jensen, Call, & Tomasello, 2012). This is interesting because the punishment is costly for them and yields no material gain, so it is also be called as altruistic punishment (E Fehr & Gächter, 2002).

However, in our life, third-parties are always involved in norm violation where the perpetrators are nevertheless strangers. Consider a following case: a judge has to make the judicial verdict on his beloved son who conducts a crime of theft. Could this judge make the just judgment, i.e., treating his son in a same way as a stranger criminal? This example introduces an important research question: whether and how does social distance (SD) modulates TPP? It has been robustly showed that SD could affect our choices in many contexts, such as that the generosity to others decreased as a function of social distance (Strombach et al., 2015). And previous findings had proved that the decreasing generosity with social distance is a robust phenomenon, with respondents across settings and cultures reliably willing to sacrifice more resources for socially close others relative to distant others (Ma, Pei, & Jin, 2015; Strombach et al., 2014). However, how social distance modulates our punishment behavior has not been studied yet. When the norm violators were close others, conniving them would break the social norm whereas punishing them would hurt close others. Regarding this, it is different from previous studies in which the punishment behaviors were always regarded as social rewards (Behrens, Hunt, & Rushworth, 2009; De Quervain et al., 2004; Ohtsuki, Iwasa, & Nowak, 2009), and it’s a really tough conflict when we have to make such a decision. Thus, whether we would connive close others or we are more severe to them, is still unknown.

Why do we punish the norm violators? Ernst Fehr and Schmidt (1999) suggested that people were averse to inequality per se, they would like to move in the direction of equitable outcomes with others by their own cost. This inequality-sensitive psychology was further proved by a lot of researches (Blake et al., 2015; Raihani & McAuliffe, 2012), Dawes and his colleagues (2007) also proved this view by observing the punishment increases with the severity of the violation. However, in third party punishment, the inequity aversion could either between self and other parties, or between the first party and second party. Based on the different inequity aversion, there are two main perspective. One is called “egocentric inequity”, defined as the absolute payoff difference between the decision-maker and other parties. That is, the inequity aversion comes from when people compare their payoff both with the proposer and the receiver (Ernst Fehr & Schmidt, 1999). Another is called “other-regarding inequity”, defined as the absolute payoff difference between the proposer and the receiver. That is, the inequity aversion comes from the unfair allocations between the two parties rather than subjects themselves (Zhong, Chark, Hsu, & Chew, 2016). However, if the norm violator is the close others, whether the social distance would modulate our inequity aversion remained to be further investigated.

On the neural level, previous studies of TPP took either the hypothetical judicial decision-making context or behavioral economic paradigm where both the proposers and the receivers were strangers to participants (Buckholtz et al., 2008; Ernst Fehr & Fischbacher, 2004). For example, a recent study found region in the mentalizing network and the central-executive network were important in TPP (Bellucci et al., 2017). The punishment behavior is generally considered to be related to the mentalizing network for evaluating legal responsibility and the central-executive network for determining appropriate punishment (Bellucci et al., 2017). The mentalizing network including the medial prefrontal cortex (MPFC) and the temporoparietal junction (TPJ), and is induced by other-regarding value, that is to say the empathy toward the victim (Batson et al., 2007; Baumgartner, Götte, Gϋgler, & Fehr, 2012; Bellucci et al., 2017), and we could consider it as the social norm. The central-executive network including the dorsolateral prefrontal cortex (dlPFC) converts the blame signal into a specific punishment decision (Krueger & Hoffman, 2016).

Regarding the effect of social distance, A recent fMRI study (Strombach et al (2015) investigated the neural underpinnings of social distance related sharing behavior, and found that the TPJ supports social distance related decisions by modulating basic neural value signals in the ventromedial prefrontal cortex (vmPFC) to incorporate social-distance-dependent other-regarding preferences, and the vmPFC valuated the signals associated with both selfish and generous choice options. Similarly, Hill, Yi, Spreng, and Diana, (2017) also found that the social distance is related to areas comprising frontoparietal control, default, and mesolimbic reward networks. Although the third party punishment and the social distance both cares about “other-regarding”, the “other-regarding” in third party punishment represents the social norm while in social distance represents close other’s benefits. Even we distinguish the two types of “other-regarding”, we could find the neural basis were similar. We infer that could because there is no conflict between the two types of “other-regarding”, that is to say, when we make a punishment, we maintained the social norm as well as close other’s benefits. However, there is no study combine the two types of “other-regarding” together and make them conflict to explore the neural mechanisms.

In the present study, we adopted a modified third-party punishment task by using fMRI to investigate how SD affects TPP both at the behavioral and the neural level. Specifically, participants in the scanner took the role of the third-party observer, who was presented with a series of monetary split advocated by a proposer to a receiver. Crucially, we only manipulated the social distance between the proposer and participants, whereas the receiver was always a stranger. Participants were asked to indicate how much money of their own they would use to punish the unfair proposer (i.e., reducing proposer’s payoff) with differential SD to them. Such that we could observe a conflict between the social norm and the other benefits. Based on previous literature, we hypothesize that people would use more money to punish the strangers than close others. Moreover, we would like to explore the neural underpinnings of the social distance related third party punishment.

## 2. Method

### 2.1 Participants

Thirty-four undergraduates (mean age = 20.39, SD = 1.46; 19 men) were recruited via online fliers. All participants were right handed and had no history of psychiatric or neurological disorders. They all gave informed consent and the procedure was approved by the ethics committee of the South China Normal University.

### 2.2 Procedure and Tasks

#### Pre-scanning phase: social distance (SD) manipulation

Once arrival, participants received verbal and written instructions for the tasks. Following the procedure by Strombach *et al* (2015), participants were first asked to rate their perceived closeness to specific persons in their social environment on a 100-point scale (i.e., mother, father, siblings, grandparents, kin, best friend, roommates, circle of friends, colleagues, neighbors, acquaintances, lover and stranger). They skipped the rating for the person who did not exist in the social environment (e.g., lover). Before entering the scanner, participants were asked to write down only one name of following social distances from their social environment: 1, 2, 3, 5, 10, and 20. Notably, we also included social distance levels of 50 and 100 in the experiment, i.e., 50 represents the person participants have once met but do not know any other information about; and 100 represents the strangers. Hence, participants were not required to indicate names for persons in these two SDs. Besides, participants were explicitly asked to only include individuals toward whom they did not have a negative attitude.

There were 2 sessions of practice before the scanning, which were same as the scanning procedure except the order of the trials, and was aimed to make participants get used to the procedure in the scanner. After completing the scanning session, participants received a 100 CYN show-up fee in the experiment.

#### Scanning-phase: the modified third-party punishment task

We adopted a modified third-party punishment (TPP) task in the current fMRI study. Participants were instructed to imagine a situation where a proposer (labeled as player A) and a receiver (labeled as player S) was involved. The proposer was endowed with 100 CNY and could freely allocate the endowment to the receiver, who had to receive the allocation. Decisions from proposers could be seen by participants in the current study. Participants, as unaffected third-party observers, could decide how much of their own endowment to punish the unfair proposer. The key manipulation here was to introduce proposers with different SDs to participants, which distinguished our design from the standard TPP game. In specific, SD was displayed in a scale consisting of 100 icons (see Figure 1C). The white icon at the left end of the scale represents themselves; the blue icon stands for a specific person A in their social environment; the number under the blue icon stands for the SD between the participants and A; the gray icon at the right end of the scale represents the receiver S, whose SD was always 100.

**Figure 1.**
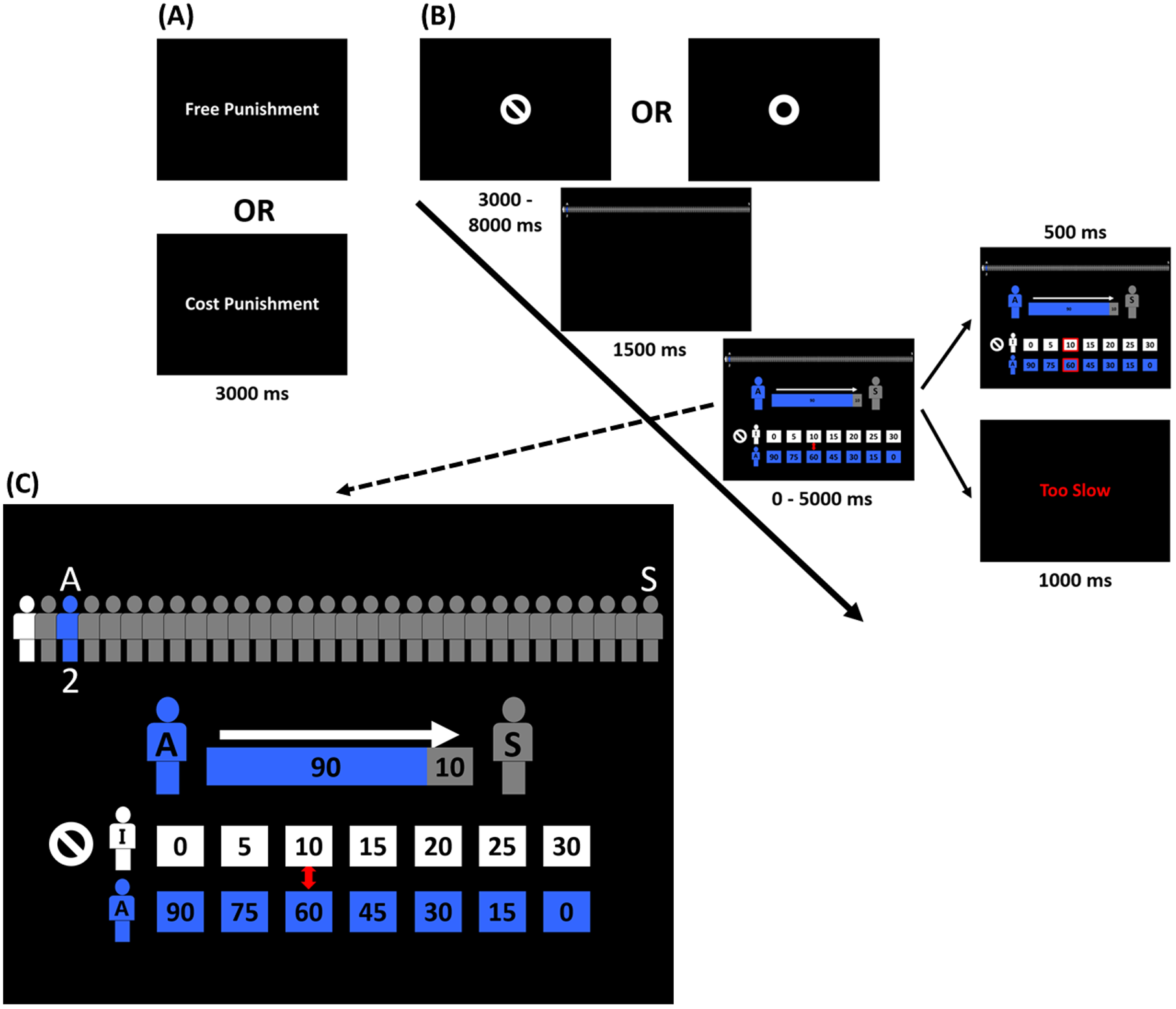
Trial procedure. **(A)** At the beginning of each block, there was an instruction screen showing the type of punishment (i.e., “free punishment” or “costly punishment”). **(B)** Each trial started with a cue (i.e., a circle means costly punishment, a circle with a line inside means free punishment). Then the social distance information for this trial was given on top of the screen. After that, the participants would be presented with the allocation and the punishment options. They needed to choose one punishment level from 7 options, i.e., 0, 5, 10, 15, 20, 25, 30, within 5000 ms. Then the selected option would be highlighted in red as a feedback. If participants failed to make the decision in 5000 ms, a warning screen would be shown. The total time of a trial was 10000 ms (jittered by ±1000 ms), and the duration for the cue was dependent on the reaction time of the previous trial. **(C)** An example display of decision phase: on the top we displayed only 11 icons, instead of 101 icons shown during scanning, to facilitate perceptibility.

The fMRI scanning included 6 sessions with each containing 54 trials. Among them, there were 48 target trials displaying unfair allocations equally distributed to 12 blocks (i.e., 4 in each block). In half blocks, participants could punish the proposer by reducing his/her payoff at the cost of their own endowment (i.e., costly condition); in the other half, they could punish the proposer without any cost to their endowment (i.e., free condition). The target trials covered all 24 combinations between social distance (i.e., 1, 2, 3, 5, 10, 20, 50, 100) and unfair allocation (i.e., 90 vs 10, 85 vs 15, 80 vs 20), with each combination appearing once for each punishment condition respectively. Besides, we added 6 filler trials displaying quasi-fair allocations (i.e., 65 vs 35, 60 vs 40, 55 vs 45) randomly assigned to 6 blocks to make the situation more natural. All blocks and trials were presented pseudo-randomly.

Each block began with a 3000ms notification of the punishment condition (see Figure 1A). In each trial, participants were endowed with 50 CNY. The trial started with a cue of the punishment type (i.e., costly or free), which lasted for a jittered interval (i.e., 3000 – 8000 ms). Next, a 1500ms-screen with the SD information of the proposer was displayed, which was followed by the decision screen. In this screen, participants saw the money allocation from the proposer to the stranger, and were provided with seven options with different punishment levels (i.e., 0, 5, 10, 15, 20, 25, 30 CNY) and the corresponding payoff for the proposer after punishment (i.e., 90, 75, 60, 45, 30, 15, 0 CNY). They needed to select one of them within 5000 ms, by pressing two buttons to move the cursor (with a random initial position) and confirming the final choice with another button with their right hand. The participants were required not to move the cursor until they determined the final option. Once they confirmed the choice, a red frame appeared on the chosen-option for 500 ms, and the rest time of decision phase (i.e., 5000ms – decision time) were added to the jitter cue. If they failed to response within 5000ms, a warning screen would be shown (see details in Figure 1B).

### 2.3 Computational Modelling

In order to investigate the computation mechanism underlying social distance (SD)-dependent third-party punishment (TPP), we created a utility function which combines both the inequality aversion (i.e., an important form of social preference) as well as the social distance. Based on previous literature, we established our models on the following well-known models: namely 1) the classical self-centric Fehr-Schmidt inequality aversion model (FSIA; Equation 1-2) (Ernst Fehr & Schmidt, 1999), 2) the other-regarding third-party inequality aversion model (TPIA; Equation 3-4) (Zhong et al., 2016), and 3) the modified hyperbolic discounting function (Equation 5-7) concerning the social distance (Jones & Rachlin, 2006). For each model, we derived two variations (i.e., FSIA_m1, FSIA_m2, TPIA_m1 & TPIA_m2) with differential assumptions on unknown parameters. Below are model details.

***FSIA_m1***:

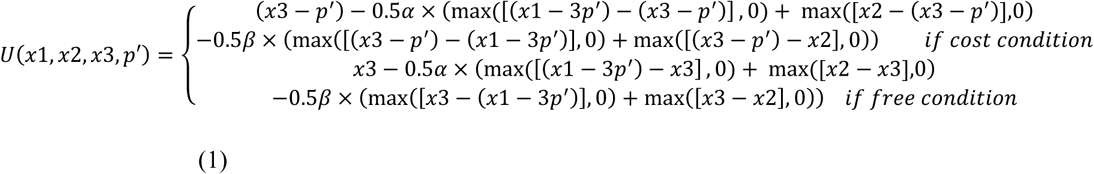

*Note*: This is the three-person version of Fehr-Schmidt model;

*U*: subjective utility of the participant (i.e., third-party decision maker) given the punishment option; x1/x2/x3: initial payoff of the Player A/Player B/the participant (i.e., x3 always equals to 30);

*p’*: punishment amount by the participant (assuming the Player A is a completely stranger for the participant)

*α*: disadvantageous inequality aversion (i.e., degree of aversion if participant gets less than others; 0 ≤ *α*≤ 5)

*β*: advantageous inequality aversion (i.e., degree of aversion if participant gets more than others; 0 ≤ *β*≤1)

***FSIA_m2***:

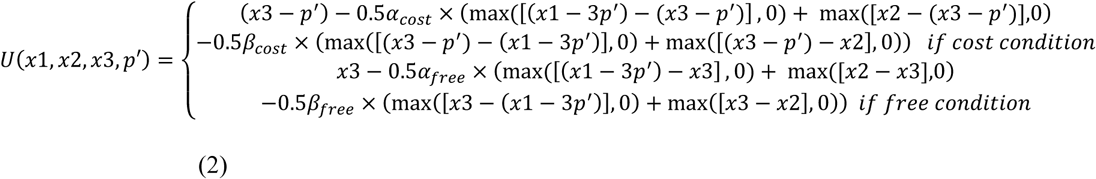

*Note*: *α*_cost_/*α*_free_: disadvantageous inequality aversion in cost/free condition;

*β*_cost_/*β*_free_: advantageous inequality aversion in cost/free condition;

The rest are the same as *FSIA_m1*.

***TPIA_m1***:

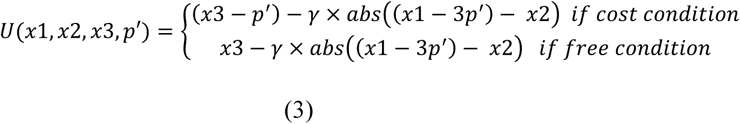

*Note*: This is the other-regarding third-party inequality aversion model;

γ: other-regarding inequality aversion (i.e., degree of aversion to the inequality between the Player A and the Player B; 0 ≤ *γ* ≤1)

***TPIA_m2***:

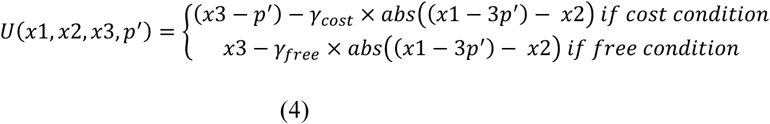

*Note*: γ_cost_/γ_free_: other-regarding inequality aversion in cost/free condition; The rest are the same as *FSIA_m1*.

To take into account the social distance, we modified the hyperbolic discount function to the following form:

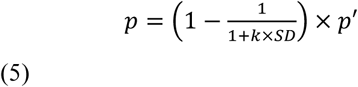

*Note*: *SD*: social distance (i.e., 1, 2, 3, 5, 10, 20, 50, 100);

*p*: actual punishment amount by the participant given the SD (i.e., 0, 5, 10, 15, 20, 25, 30);

*k*: discount rate (i.e., how fast the punishment amount discounts with closer SD; 0 < *k* ≤ 1); The rest are the same as *FSIA_m1*.

Thus, the equation could be re-formulized as the following:

For *FSIA_m1* & *TPIA_m1*:

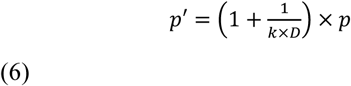

*Note*: *p’/SD/p/k*: same as Equation 5.

For *FSIA_m2* & *TPIA_m2*:

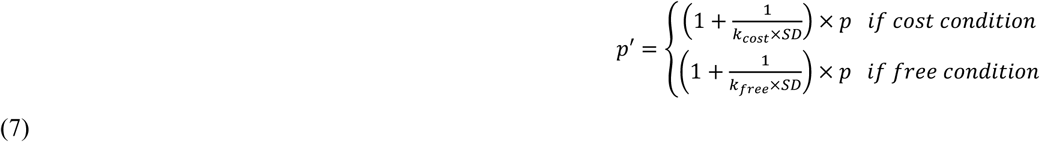

*Note*: *k*_*cost*_*/k*_*free*_: discount rate in cost/free condition;

The rest are the same as Equation 5.

By using softmax function (equation 4), we could then compute the probability of the subjective utility of the chosen option (*Uc*; we will treat this as a categorical logit problem, namely choosing one out of seven options; *p* could be 0, 5, 10, 15, 20, 25, 30):

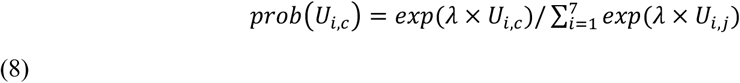

*Note*: *U*_*i,c*_: subjective utility of the participant (i.e., third-party decision maker) of the chosen options *c* in each trial *i*;

*λ*: the inverse temperature parameter which denotes the sensitivity of a participant’s choice to the difference in utility between the left and right option (0 ≤ *λ* ≤ 5)

Model estimation and comparison were performed by Rstan (http://mc-stan.org/interfaces/rstan) and relevant R packages. By using hierarchical Bayesian modelling, a state-of-art modelling technique which considers individual and group parameters (i.e., posterior distribution) in a simoutaneous and muturally constraining fashion during estimation (Ahn, Haines, & Zhang, 2016), we can compare which model fits our data the best and then estimate parameters which capture the social preference (e.g., inequality aversion) for each individual.

Moreover, we extracted the estimated parameters from the winning model to compute the trial-wise utility given the chosen option; these values were then used as the parametric modulator in the fMRI analyses so that we could test which brain areas track the computation during decision making (O’doherty, Hampton, & Kim, 2007).

### 2.4 Data Acquisition and Preprocessing

Scanning was performed on a 3-tesla Trio Scanner (Siemens). Functional data were acquired using echo-planar imaging sequences (repetition time = 2 s, echo time = 30 ms, flip angle = 90°, field of view = 224 mm, slice thickness = 3.5 mm). A total of 32 axial slices were acquired in interleaved order (in-plane resolution 3 × 3 mm). Anatomical images were T1-weighted (MDEFT, 1 × 1 × 1 mm resolution). The presentation of the task and recording of behavioral responses were performed using E-Prime software, version 2.0. Neural data of 34 participants were analyzed using SPM12 (http://www.fil.ion.ucl.ac.uk/spm/software/spm12/). The results are visualized using the xjview toolbox. Two participants were excluded due to excessive head movements during scanning (>2 mm translation or >2°rotation), and four participants had to be excluded one session of the data due to head movements.

Functional images were realigned using a six-parameter rigid-body transformation. Each individual’s structural T1 image was co-registered to the average of the motion-corrected images using 12-parameter affine transformation. Individual T1 structural images were segmented into grey matter, white matter, and cerebrospinal fluid. Functional images were, in order, slice timing correction, motion correction, segmentation using the T1-weighted image, normalized to MNI space, and smoothed with an 8mm isotropic Gaussian kernel.

### 2.5 GLM Analysis

We constructed three GLMs to identify regions responding at decision onset to the trial-wise subjective utility (GLM1), social distance (GLM2), and punishment type (GLM3), respectively.

GLM1 contained two main event regressors describing the onsets of (1) Free punishment trials;(2) Cost punishment trials. Both events were modeled as stick functions with duration zero and were each associated with a parametric modulator of the trial-wise subjective utility, and conducted an one-sample T test on second level. The key contrast we built in GLM1 was the *subjective utility*.

GLM2 contained two main event regressors describing the onsets of (1) Close distance trials; (2) Distant distance trials. Both events were modeled as stick functions with duration zero and using social distance sensitivity (ie., punishment difference of close distance – distant distance, that means if the larger social distance sensitivity is, the more someone distinguishes close distance and distant distance.) as a covariate on second level. The key contrast was *close distance – distant distance*.

GLM3 contained two main event regressors describing the onsets of (1) Free punishment trials; (2) Cost punishment trials. Both events were modeled as stick functions with duration zero and using self-benefits sensitivity (ie., punishment difference of free condition – costly condition, that means if someone distinguishes free and costly condition more clearly, the more he cares about his own payoff, because the only difference between free and costly condition is whether the participants need to consume his own payoff) as a covariate on second level (Table 1A). The key contrast was *free condition – costly condition*.

**Table 1.** GLM info. We have 3 GLMs in total, all of them contained two main events and six events of no-interest. And all the events were stick function. **(A)** The details of the three GLM; **(B)** the details of the six events of no-interest.

| GLM | Main Event | Parametric | Onset | Second Level |
| --- | --- | --- | --- | --- |
| GLM1 | 1) Free punishment trials | Subjective Utility | Decision phase | One-sample T test |
|  | 2) Costly punishment trials | Subjective Utility | Decision phase |  |
| GLM2 | 1) Close distance trials (SD = 1, 2, 3, 5) | None | Decision phase | Add covariate: Social distance sensitivity |
|  | 2) Distant distance trials (SD = 10, 20, 50, 100) | None | Decision phase |  |
| GLM3 | 1) Free punishment trials | None | Decision phase | Add covariate: Self-benefits sensitivity |
|  | 2) Costly punishment trials | None | Decision phase |  |

**(B)**
| Onset of Event for No Interest | Parametric |
| --- | --- |
| 1) The verbal instruction in the beginning of each block | None |
| 2) The social distance information of each trial | None |
| 3) The punishment options for filler trials | None |
| 4) The punishment options for no-responded trials | None |
| 5) The feedback for responded trials | None |
| 6) The feedback for no-responded trials | None |

All three GLMs also contained six additional event regressors of no interest, describing the onsets of: (1) The verbal instruction in the beginning of each block; (2) The social distance information of each trial; (3) The punishment options for filler trials; (4) The punishment options for no-responded trials; (5) The feedback for responded trials; (6) The feedback for no-responded trials. These events were all modeled as stick functions with duration zero. Finally, six motion regressors obtained during realignment were included to control for motion of no interest (Table 1B). All images were high-pass filtered in the temporal domain (filter width 128 s). Autocorrelation of the hemodynamic responses was modeled as an AR(1) process.

Random effects models were done in SPM12 by specifying a separate general linear model for each participant and pooled at the second level.

### 2.6 Functional connectivity analyses

To investigate the functional connectivity between vmPFC and other regions in the brain, we conducted a seed-to-voxel general psychophysiological interaction (gPPI) analysis by CONN (http://www.nitrc.org/projects/conn). We created vmPFC seed regressors by using a 6-mm spheres surrounding the peak voxels of vmPFC in GLM3 contrast *close distance – distant distance*, showing negative effects resulting from facing close distance relative to distant distance. Six motion regressors obtained during realignment were added to the PPI GLM to control for the effect driven by head motion.

For small-volume correction analysis, we use coordinates from a meta-analysis study of ACC, vmPFC, and ventral striatum (Bartra, Mcguire, & Kable, 2013), vmPFC and ventral striatum in this paper were related to the subjective value effected by monetary incentives while ACC was related to the subjective value effected by positive effects in the reward domain, all of them were based on MNI coordinates. More specifically, we used coordinates of ACC (x = −2, y = 24, z = 26) with a 20-mm sphere (Onur, Piefke, Lie, & Thiel, 2011), vmPFC (x = 0, y = 52, z = −8) with a 15-mm sphere (Cheng et al., 2015) and with ventral striatum (x = 14, y = 14, z = −6) a 10-mm sphere (Hermans et al., 2010). We also conducted a small volume correction for left hippocampus by aal templates. All these SVC results passed a corrected significance threshold of p < 0.05.

## 3. Results

### 3.1 Behavior Results

#### Punishment level

We conducted a repeated-measure ANOVA with the punishment amount as the dependent variable, type of punishment (costly / free) and social distance (SD; 1, 2, 3, 5, 10, 20, 50 and 100) as within-subject factors. We found that participants punished the proposer more severely when it is free (vs. cost; *F* (1, 32) = 30.65, 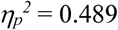, *p* < 0.001) and when their SD increased (*F* (7, 224) = 47.20,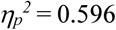, *p* < 0.001). Interestingly we observed the interaction between punishment type and social distance (*F* (7, 224) = 16.93,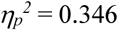, *p* < 0.001). Post-hoc analyses revealed that the lager SD was, the lager difference was shown in punishment between cost and free condition (see Figure 2).

**Figure 2.**
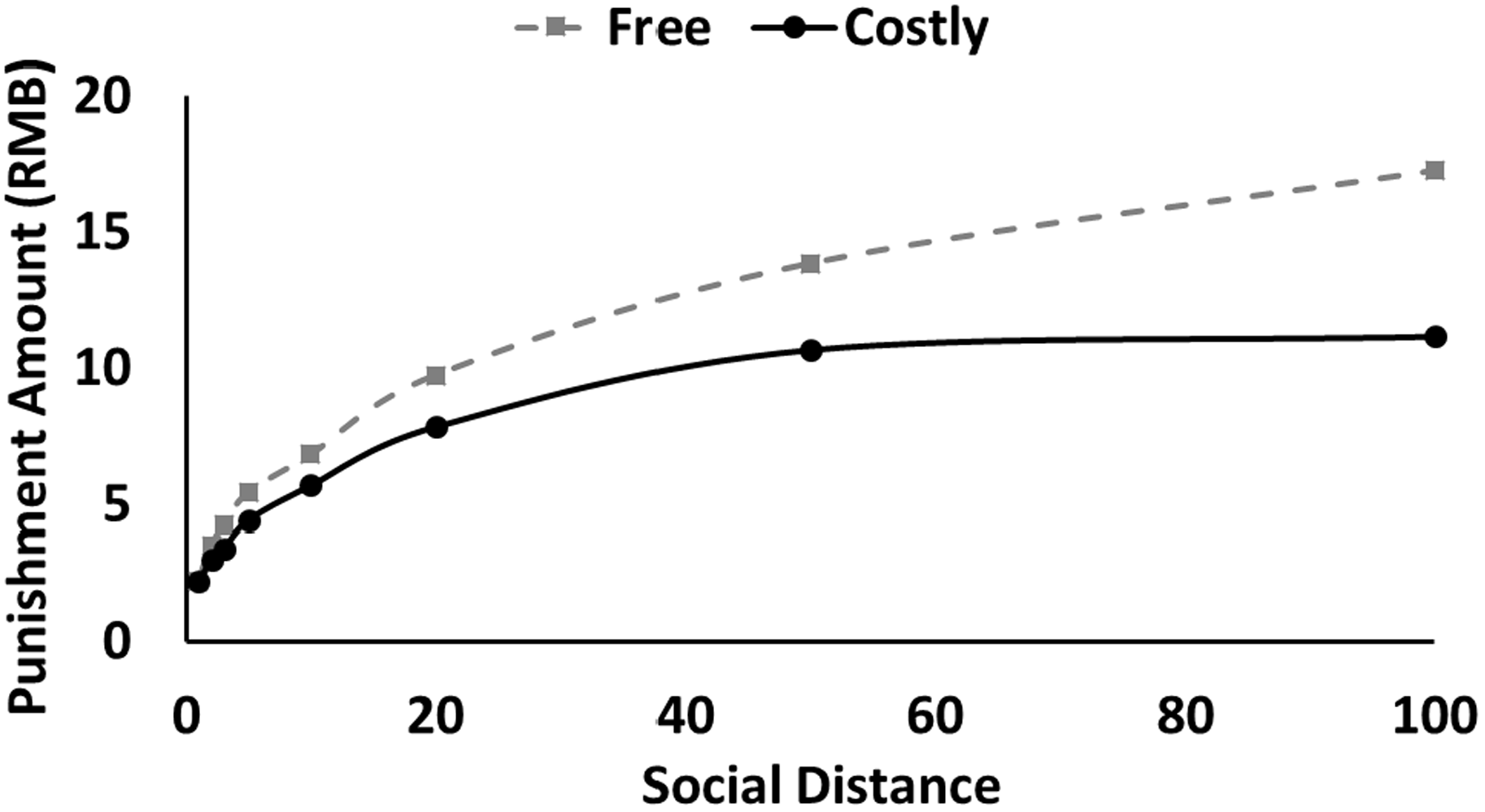
Results of punishment amount. The punishment amount changed as a function of both punishment condition and SD.

**Figure 3.**
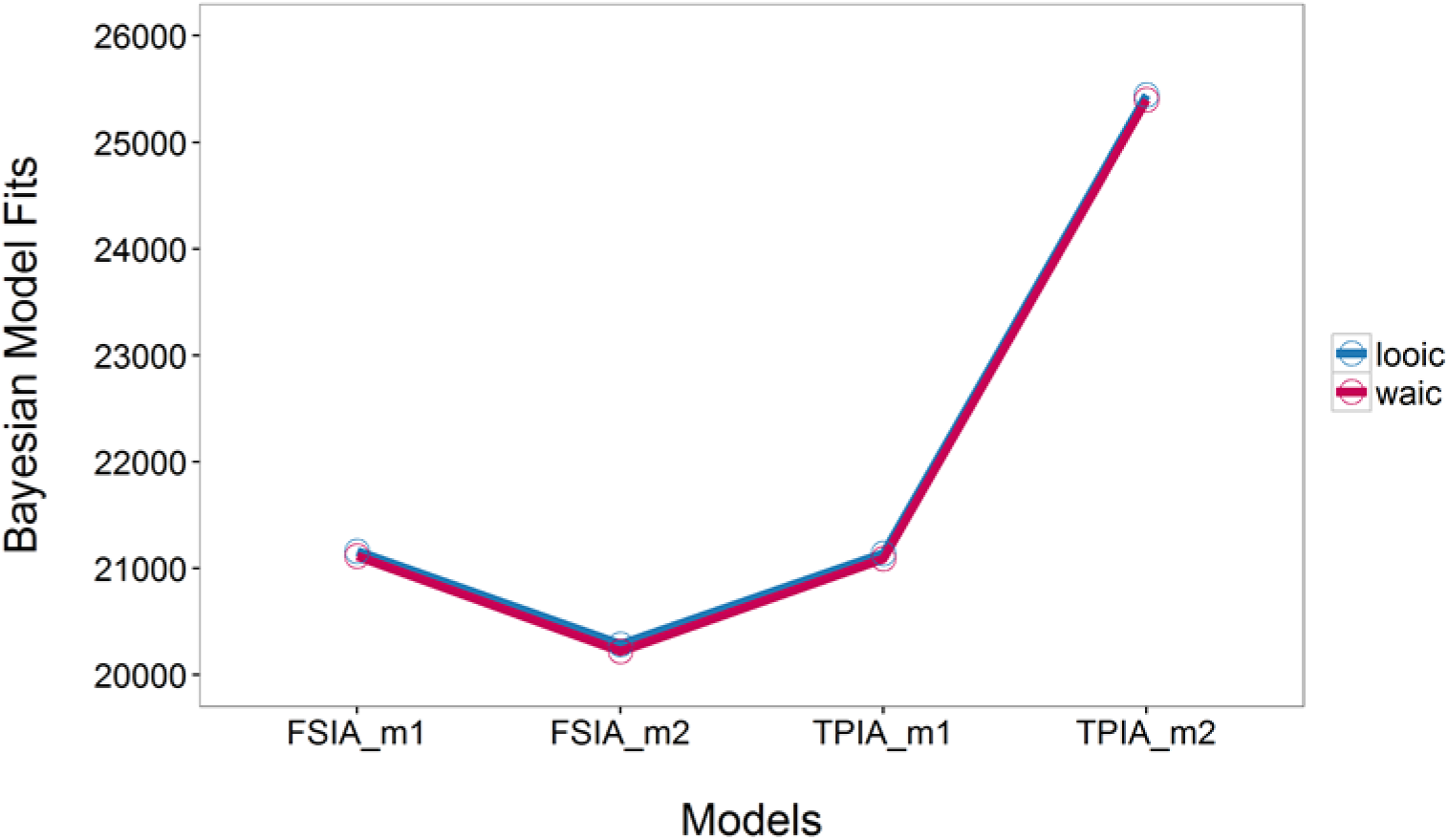
Bayesian Model Comparison. Lower looic and WAIC score indicates better out-of-sample prediction accuracy of the candidate model. Looic refers to leave-one-out information criterion, and WAIC refers to widely applicable information criterion.

#### Reaction time

One participant was excluded from our data analysis because of random choice. First we conducted a repeated-measure ANOVA with the reaction time of each trial as the dependent variable, type of punishment (costly / free) and social distance (SD; 1, 2, 3, 5, 10, 20, 50 and 100) as within-subject factors. We found that participants spend more time to make a decision when the SD increased (*F* (7, 224) = 14.98, 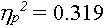, *p* < 0.001), but the difference on reaction time between cost and free conditions only appears when SD is 10 (*p* = 0.003). The results were after Bonferroni correction.

#### Bayesian Model Comparison

The hierarchical Bayesian model comparison showed that the second model of Fehr-Schmidt inequality aversion model (i.e., FSIA_m2) which distinguished advantageous/disadvantageous inequality aversion parameters in terms of cost/free condition was with the lowest the leave-one-out information criterion (LOOIC) and widely applicable information criterion (WAIC) scores. Lower LOOIC or WAIC scores indicate better out-of-sample prediction accuracy of the candidate model (Vehtari, Gelman, & Gabry, 2016). This result suggested that FSIA_m2 outperformed the rest competitive models (see figure 2).

### 3.2 fMRI Results

Firstly, we searched for brain regions encoding subjective utility (parametric modulator, GLM1) during decision phase. The results showed that with a lower subjective utility, right dlPFC were more engaged (please see Figure 4A). Then we extract the beta value of right dlPFC using a 6-mm sphere based on the peak coordinates, to see how it changes in different punishment type and social distance, and it revealed that the neural activation of right dlPFC had the same pattern as behavior results.

**Figure 4.**
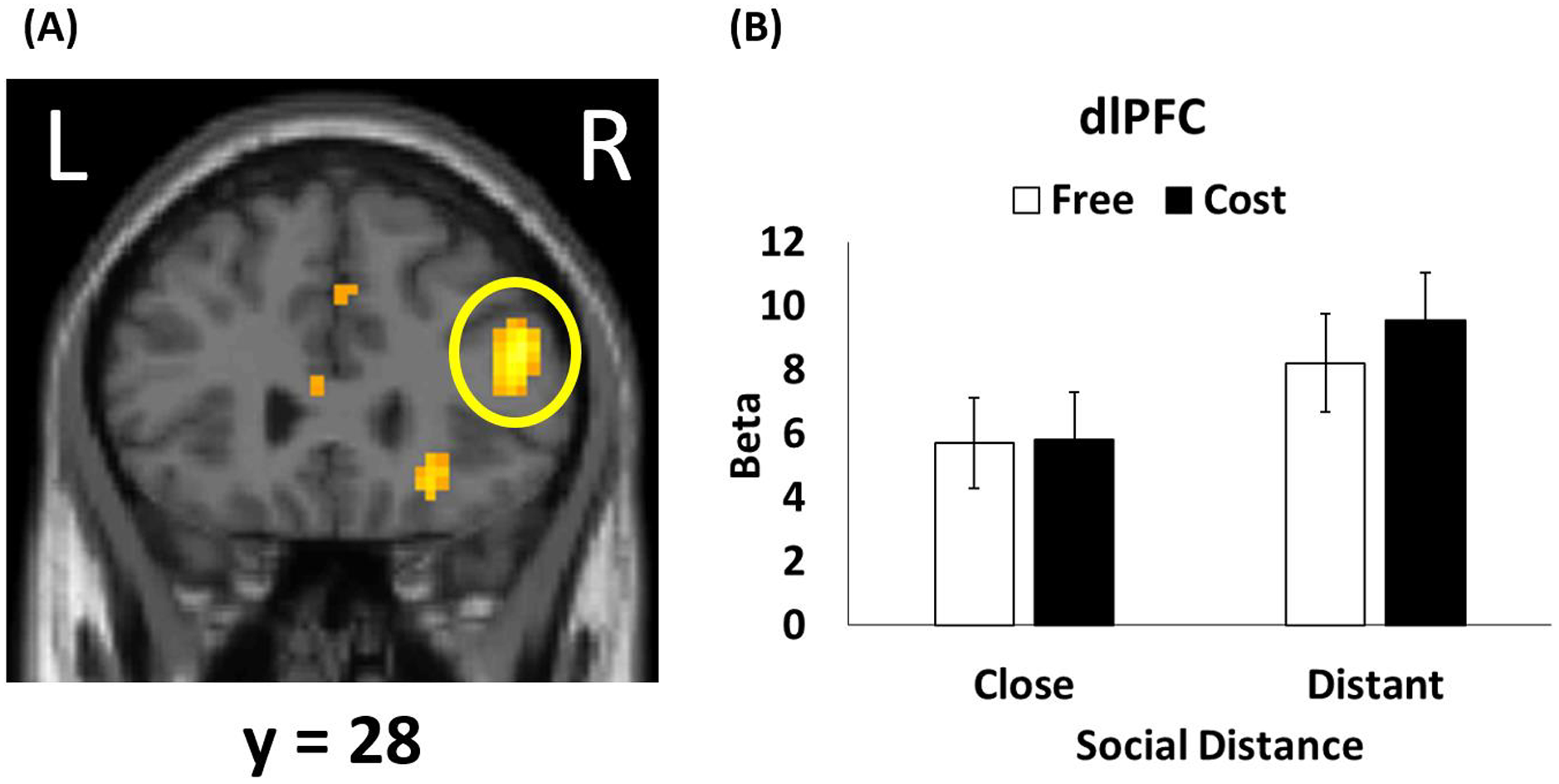
Model-based fMRI results (GLM1). **(A)** Right dlPFC (peak MNI coordinates 54, 18, 24; t(30) = 6,88, P_FWE_ < 0.001) negative correlated with subjective utility of the chosen option based on the winning model. **(B)** The beta value of the peak activation of right dlPFC changed as a function of both punishment types and SD. The error bar refers to SEM.

Then we explored the neural activation related to social distance in GLM2, which is the core factor in our study. We used the “social distance sensitivity” as a second level covariate, and observed a reduced activation in the vmPFC and bilateral hippocampus (vmPFC and left hippocampus were small volume corrected) of the contrast *close > distant* (see Figure 5A). We also plot scatters to see the correlation between the social distance sensitivity and the corresponding beta values for each ROI (see Figure 5B), as well as the correlation between the punishment amount and the corresponding beta values for each ROI (see Figure 5C).

**Figure 5.**
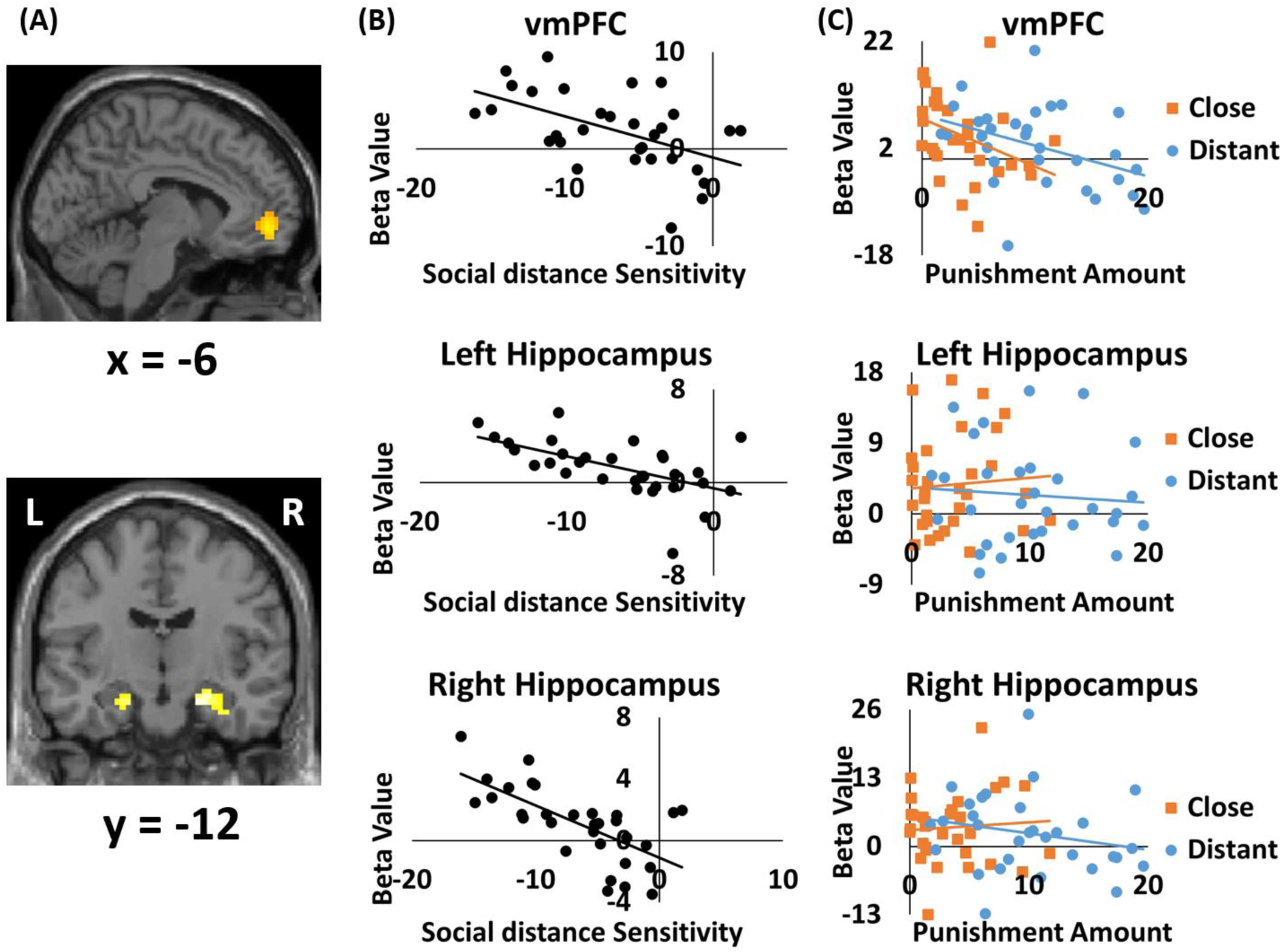
Modulation of SD on the relationship between neural activation and punishment level (GLM2). **(A)** Differential neural activation in vmPFC (Peak MNI coordinates −6, 51, −6; t(29) = 3.69, P_SVC-FWE_ = 0.050) and bilateral hippocampus (Peak MNI coordinates for right hippocampus 24, −15, −21; t(29)= 5,06, P_FWE_ = 0.024; for left hippocampus −21, −12, −21; t(29) = 4.01, P_SVC-FWE_ = 0.04) negatively correlated with the social distance sensitivity (i.e., close distance vs. distant distance). **(B)** Scatter plot depicting the correlation between the social distance sensitivity and the differential beta values (i.e., close vs. distant) in corresponding ROIs (spheres with a radium of 6-mm centering at the local peak MNI coordinates) This figure is only for display and no statistical analysis was performed. **(C)** The correlation between the respective punishment amount (close & distant) and the beta value of the corresponding ROI. close: trials with SD = 1, 2, 3, 5; distant: trials with SD = 10, 20, 50, 100.

Furthermore, functional connectivity analysis by gPPI approach revealed that the left dlPFC demonstrated enhanced connectivity with the vmPFC in close distance compared to distant distance during decision-making period (see Figure 6).

**Figure 6.**
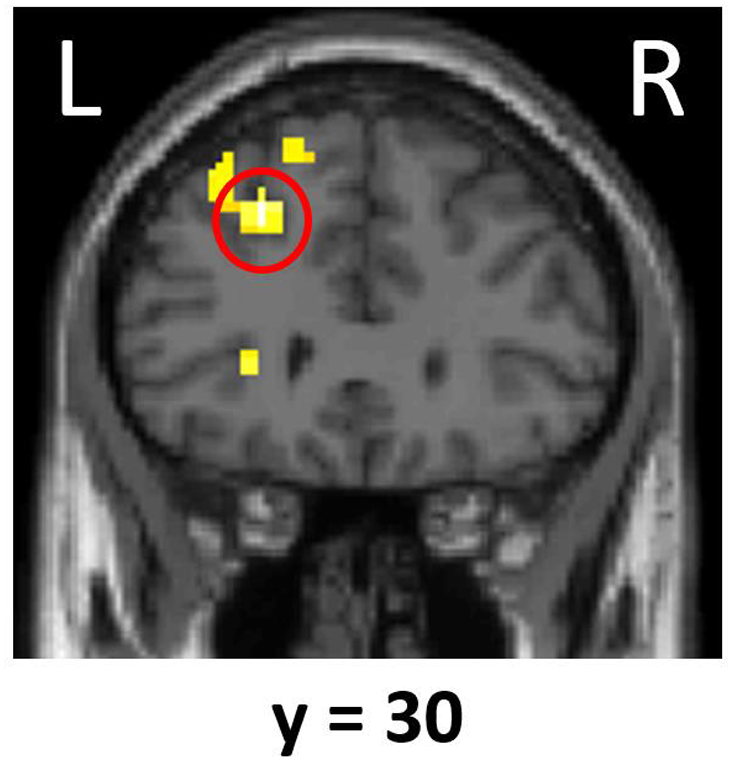
Functional connectivity results (GLM2). The left dlPFC. (peak MNI coordinates −24, 30, 42; *t*(30)= 5.65, *P*FWE = 0.001) **displays higher** the functional connectivity with the vmPFC (seed region; peak MNI coordinates −6, 51, −6) during decision-making period when the allocator is close (vs. distant) to participants.. close: trials with SD = 1,2,3,5; distant: trials with SD = 10,20,50,100.

Finally, as the computational model suggested that we had different inequity aversion in free and costly condition, the two punishment types were compared in GLM3. We define the punishment difference between free and costly condition (ie., punishment in free condition – punishment in costly condition) as “self-benefits sensitivity”, that means if someone distinguishes free and costly condition more clearly, the more he cares about his own payoff, because the only difference between free and costly condition is whether the participants need to consume his own payoff. We use this “self-benefits sensitivity” as a second level covariate, and observed a reduced activation in the dorsal anterior cingulate cortex (dACC) and ventral striatum (both small volume corrected) of the contrast *free > costly* (see Figure 7A). We also plot scatters to see the correlation between the self-benefits sensitivity and the corresponding beta values for each ROI (see Figure 7B), as well as the correlation between the punishment amount and the corresponding beta values for each ROI (see Figure 7C).

**Figure 7.**
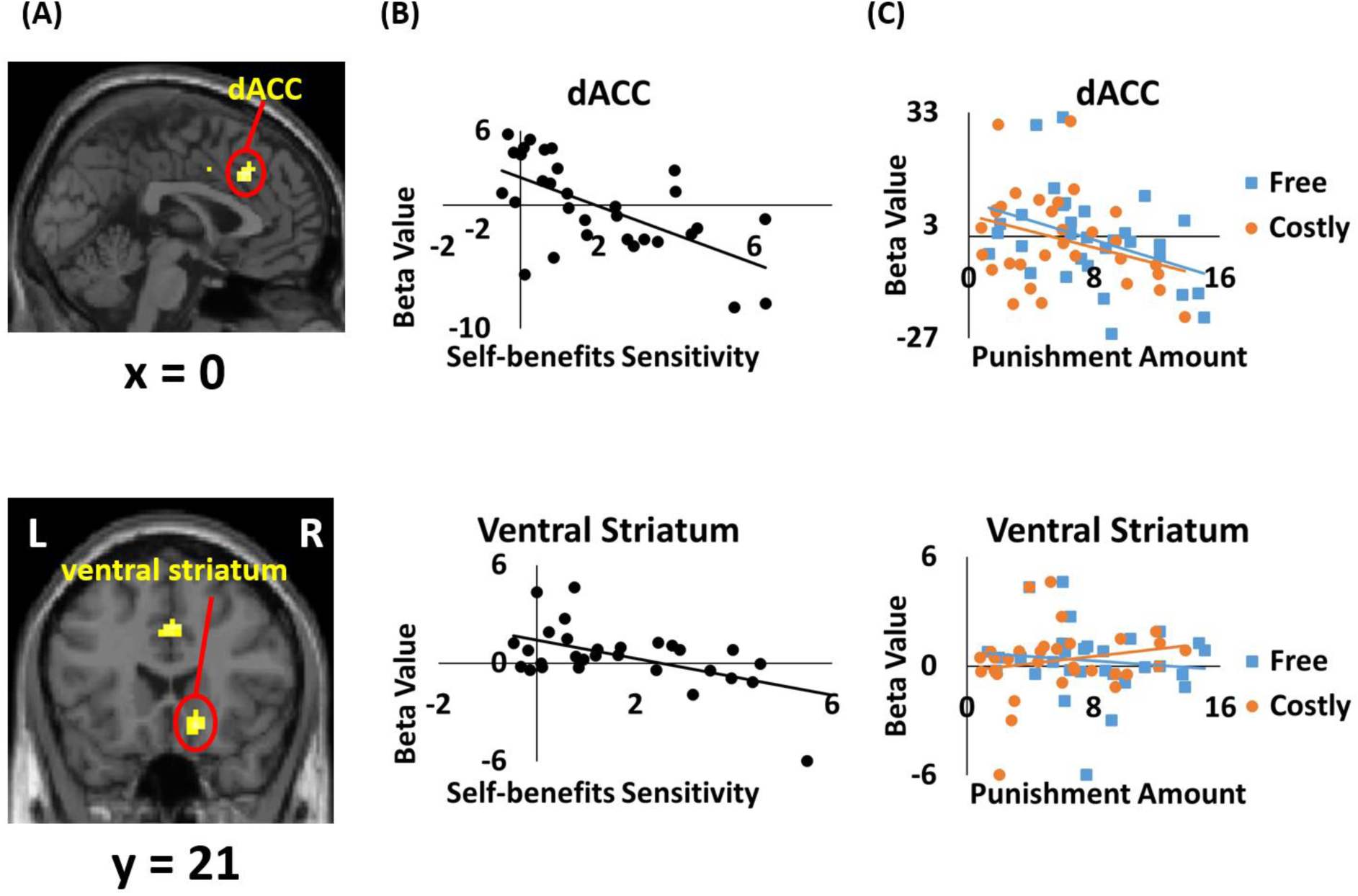
Modulation of punishment type on the relationship between neural activation and punishment level (GLM3). **(A)** We found the differential neural activation in dACC (peak MNI coordinates 0, 27, 36; t(29) = 4.48, P_SVC-FWE_ = 0.018) and ventral striatum (peak MNI coordinates 15, 21, −12; t(29) = 4.25, P_SVC-FWE_ = 0.005) were negative correlated with the self-benefits sensitivity (i.e., free vs. costly). **(B)** Scatter plot depicting the correlation between the self-benefits sensitivity and the differential beta value (i.e., free vs. costly) in corresponding ROIs (spheres with a radium of 6-mm centering at the local peak MNI coordinates). This figure is only for display and no statistical analysis was performed. **(C)** The correlation between the respective punishment amount (free & costly) and the beta value of the corresponding ROI.

## 4 Discussion

Norm development is so important to function well in our society, so human evolve to maintain social norms even when they were unaffected bystanders. The present study explored how social distance modulated our punishment behavior as a third party and its underlying neural mechanism. Behaviorally, we found that like previous studies which focus on generosity in different social distances (Strombach et al., 2014; Strombach et al., 2015), the punishment decisions under different social distances exhibit similar pattern, as we will connive close others in both situation. At the neural level, we found that the vmPFC and hippocampus is related to the social distance sensitivity, while the dlPFC is an integrative region which computed the subjective utility. We also observed a functional connectivity between the vmPFC and the dlPFC, indicating the two regions work together to make the final decision. Moreover, we found the dACC and the ventral striatum is associated with self-benefits sensitivity, indicating these two regions are reward-related regions.

At behavior level, the results suggested that when the close others made an unfair allocation, we were less likely to punish them. This is consistent to previous findings, as the generosity decreases as a function of social distance (Strombach et al., 2014; Strombach et al., 2015). Charness and Gneezy (2008) considered the effect of another form of social distance, they found that compared to a totally stranger, when participants knew the family name of their counterparts, they allocated a significantly larger portion of the pie. Some third party punishment studies distinguishing ingroup and outgroup also provide evidence of this view, they found that outgroup perpetrators were punished more strongly than ingroup perpetrators (Baumgartner et al., 2012; Schiller, Baumgartner, & Knoch, 2014). And it is unsurprising that participants tend to give less punishment in the costly condition, suggested that there is a cost-benefits trade-off when people making an altruistic punishment. The computational model provided us new sight about the underlying mechanism of third party punishment. As we compared the Fehr-Schmidt inequality aversion model and the other-regarding third-party inequality aversion model with the same or different inequity aversion in free and costly punishment condition respectively, and found that the FSIA model with different inequity aversion in free and costly punishment condition fits our data best. Revealed that in the present task, participants were egocentric, they mostly focus on the payoff between other parties and themselves, suggesting the possible mental process when we were making a punishment decision as a third party.

Then we explored the neural basis of the computational model combined with the fMRI analysis, it revealed that the right dlPFC was related to the subjective utility of each punishment option, while further analysis suggested that the vmPFC is related to social distance. This is consistent with previous findings that the dlPFC could integrate the information to make a decision (Domenech, Redouté, Koechlin, & Dreher, 2017; Philiastides, Auksztulewicz, Heekeren, & Blankenburg, 2011). Nejati, Salehinejad, and Nitsche (2017) found that DLPFC was both associated with cognitive and affective or reward-related executive functions. Todd A. Hare, Hakimi, and Rangel (2014) also found that DLPFC could modulate the activity in other regions and represented the weight for different attributes. In the current study, participants would face a conflict between the social norm and the close other’s benefits, and it is a complex process while they were thinking how to make the decision. On the other hand, we could observe the vmPFC is activated during tasks involving self-knowledge and person perception (Amodio & Frith, 2006; Chavez, Heatherton, & Wagner, 2016; Wagner, Haxby, & Heatherton, 2012). As numerous studies proved that the vmPFC is sensitive to self-related information, Zhu and Han (2008) found that unlike Westerners who employed the vmPFC to represent only the individual self, Chinese utilized the same brain area to represent both self and close others. And consistent conclusions were proved by Wang, Peng, Chechlacz, Humphreys, and Sui (2017) that interdependent self-construal inversely predicts the volume in the vmPFC, as interdependent self-construal usually be seen in Easterners. Similarly, a study in adolescents found that a greater overlap in neural networks for responses to self-and friend-related judgments compared to teachers and politicians (Romund et al., 2017), which is consistent with the present study that we divided the eight social distances into two categories. Interestingly, previous studies in adults indicated a stronger vmPFC involvement for stimuli that are closer or more important to oneself (Krienen, Tu, & Buckner, 2010; Murray, Schaer, & Debbané, 2012; van der Meer, Costafreda, Aleman, & David, 2010), supporting that the vmPFC is sensitive to social distance.

Unexpectedly, we did not observe TPJ neither with subjective utility nor social distance, we infer that could because in the current experiment, participants focus on the conflict between social norm and close other’s benefits rather than the feelings of other parties. But interestingly, the explorative functional connectivity analysis showed that the vmPFC showed stronger association with left dlPFC when participants punished close others compared to distant others. According to previous findings, the vmPFC supports the integration of both valuation processes and self-referential thought (D’Argembeau, 2013). Todd A. Hare et al (2014) proposed a model that the vmPFC computes the value of options by first assessing their various attributes, and then integrating them into a net value for the option as a whole. Importantly, “basic” attributes might always be represented in the final value. However, more abstract attributes are only represented in the dlPFC and modulates activity in vmPFC so that its value computations incorporate them. Many other researches supported this view that the vmPFC calculates the basic information and the dlPFC make the integration (Basten, Biele, Heekeren, & Fiebach, 2010; Domenech et al., 2017; Todd A Hare, Schultz, Camerer, O’Doherty, & Rangel, 2011). In the present study, we propose that the vmPFC process the social distance information, and the dlPFC modulates the activity in vmPFC to make the final decision. When the close others make an unfair allocation, it is a more complex process as we need to consider three aspects of conflicts: The relationship to close others, the social norm, and our own payoff. That means the dlPFC engage more in control function and we could observe a stronger functional connectivity.

Other than the vmPFC, we also observed the hippocampus associated with social distance. The traditional view of the hippocampus is that it creates a cognitive map to navigate physical space (Eichenbaum, 2015; Tolman, 1948). Tavares et al (2015) extended the cognitive maps in the hippocampus to social space. Consistent with this finding, Kumaran, Banino, Blundell, Hassabis, and Dayan (2016) also found that except mapping physical space, the hippocampus also plays an important role in charting and navigating the social distances in people’s community life. In other word, social relations may occupy a key portion of ‘memory space’, and the hippocampus worked as a hub for social navigation. Specifically, recent theory suggests that memory processes through hippocampal system are necessary to navigate social distance and social hierarchies (Laurita & Spreng, 2017). As there is not much evidence so far to draw the conclusion that the hippocampus carries a special function for social cognition, we provide new evidence regarding how the hippocampus is recruited in social contexts.

Finally, as we obtained from the inequity aversion model, people cared more about their own payoff rather than the inequity between other parties, and they had different inequity aversion in free and costly condition. Consistent with the computational model, we observed the punishment difference between free and costly conditions at behavioral level. Correspondingly, we found that the dACC and ventral striatum were associated with self-benefits sensitivity. According to previous findings, the ACC is thought to play a key role in detection of cognitive conflict (Ernst Fehr & Rockenbach, 2004) and conflict monitoring (Botvinick, Braver, Barch, Carter, & Cohen, 2001). Activity in this region is consistent with the existence of a tradeoff between self-interest and pro-social motives (Ernst Fehr & Camerer, 2007). Similar results were found in ventral striatum. Prior studies revealed that the ventral striatum exhibited neural activation during reward anticipation (Dreher, Kohn, Kolachana, Weinberger, & Berman, 2009; Yacubian et al., 2007). Further, the ventral striatum seems to be important for human cooperation and the punishment of norm violations, and it belongs to the reward-related circuits (Ernst Fehr & Rockenbach, 2004). In a word, the main function of the dACC and ventral striatum under the paradigm of third-party punishment task, was to detect and computing the cognitive conflict and cognitive value of the choices. However, the finding is not so robust in the present study as it was after the small volume correction. We suppose it could because the free condition and costly condition were both in the punishment pattern, but differed in the magnitude of inequity aversion. As we can see the beta value at dlPFC in GLM1 (Figure 4B), it indeed showed the difference between free and costly condition, but it was not significant. De Quervain et al (2004) observed similar results as they did not find significant difference between free and costly punishment. Anyway, the neural results were sufficient to support our behavior findings.

In sum, the present study revealed the underlying mechanism of social distance related third party punishment using an fMRI design combined with a computational model. Not only the mental mechanism, but also the neural mechanism. Suggesting that social distance is rather crucial to our decision making, and compared to inequity of other parties, people consider more about themselves. Future studies could further consider the neural mechanism of the inequity aversion to the first party and the second party. More important, we found the dlPFC-vmPFC connectivity in decision making, proposing that the vmPFC process the social distance information, while the dlPFC integrates other attributes like inequity and own payoff, then finally make the decision. Future studies could induce the TMS or tDCS techniques to explore whether there is a causal relationship. Moreover, we also proved that hippocampus could be associated with social navigation, not only physical navigation. This is interesting to extend the function of hippocampus, and future studies could focus more on the hippocampus in social context.

